# Auditory Corticothalamic Neurons are Recruited by Motor Preparatory Inputs

**DOI:** 10.1101/2020.05.28.121459

**Authors:** Kameron K. Clayton, Ross S. Williamson, Kenneth E. Hancock, Troy Hackett, Daniel B Polley

## Abstract

Optogenetic activation of *Ntsr1*+ layer 6 corticothalamic (L6 CT) neurons modulates thalamocortical sensory processing and perception for hundreds of milliseconds following laser offset. Naturally occurring sources of extrasensory inputs that could recruit L6 CTs prior to upcoming sensory stimuli have not been identified. Here, we found that 100% of L6 CTs in mouse primary auditory cortex (A1) expressed FoxP2, a protein marker found in brain areas that coordinate sensory inputs with movement. To test the idea that motor preparatory inputs could be a natural extrasensory activator of L6 CTs, we combined quantitative videography, optogenetically targeted single unit recordings, and two-photon imaging during self-initiated behavior. We found that A1 L6 CTs were activated hundreds of milliseconds prior to orofacial movements, but not whole-body movements associated with locomotion. These findings identify new local circuit arrangements for routing motor corollary discharge into A1 and suggest new roles for CT neurons in active sensing.

---

L6 CTs are the largest component of the corticofugal projection system and one of the largest classes of projection neurons in the brain, but their contribution to brain function and behavior remain an unsolved mystery. In the auditory cortex (ACtx), L6 CT axons bifurcate into a subcortical branch that deposits axon collaterals onto GABAergic neurons in the thalamic reticular nucleus en route to the medial geniculate body (MGB) and a vertically oriented intracortical branch that collateralizes extensively within the local vertical column, predominantly synapsing onto neurons in layers 5a and 6 (Cai et al., 2018; Guo et al., 2017; Llano and Sherman, 2008; Prieto and Winer, 1999; Winer et al., 2001) (**Fig. 1A**). Because they are spatially intermingled with other L6 cell types, and because of the strong reciprocal connectivity of the thalamus and cortex, traditional methods for *in vivo* neural recordings, stimulation or inactivation have made it challenging to identify a specific role for L6 CTs in sensory processing. The advent of optogenetic approaches to activate and silence genetically targeted L6 CT neurons in Ntsr1-Cre transgenic mice has reinvigorated research on CT circuits, inspiring new hypotheses about their role in sensory gain control and predictive coding (Crandall et al., 2015; Gong et al., 2007; Guo et al., 2017; Olsen et al., 2012; Vélez-Fort et al., 2014; Voigts et al., 2019; Williamson and Polley, 2019). However, any role for L6 CTs in active sensing has been purely speculative, as targeted recordings from L6 CTs have never been made in behaving animals.

**Figure 1.**
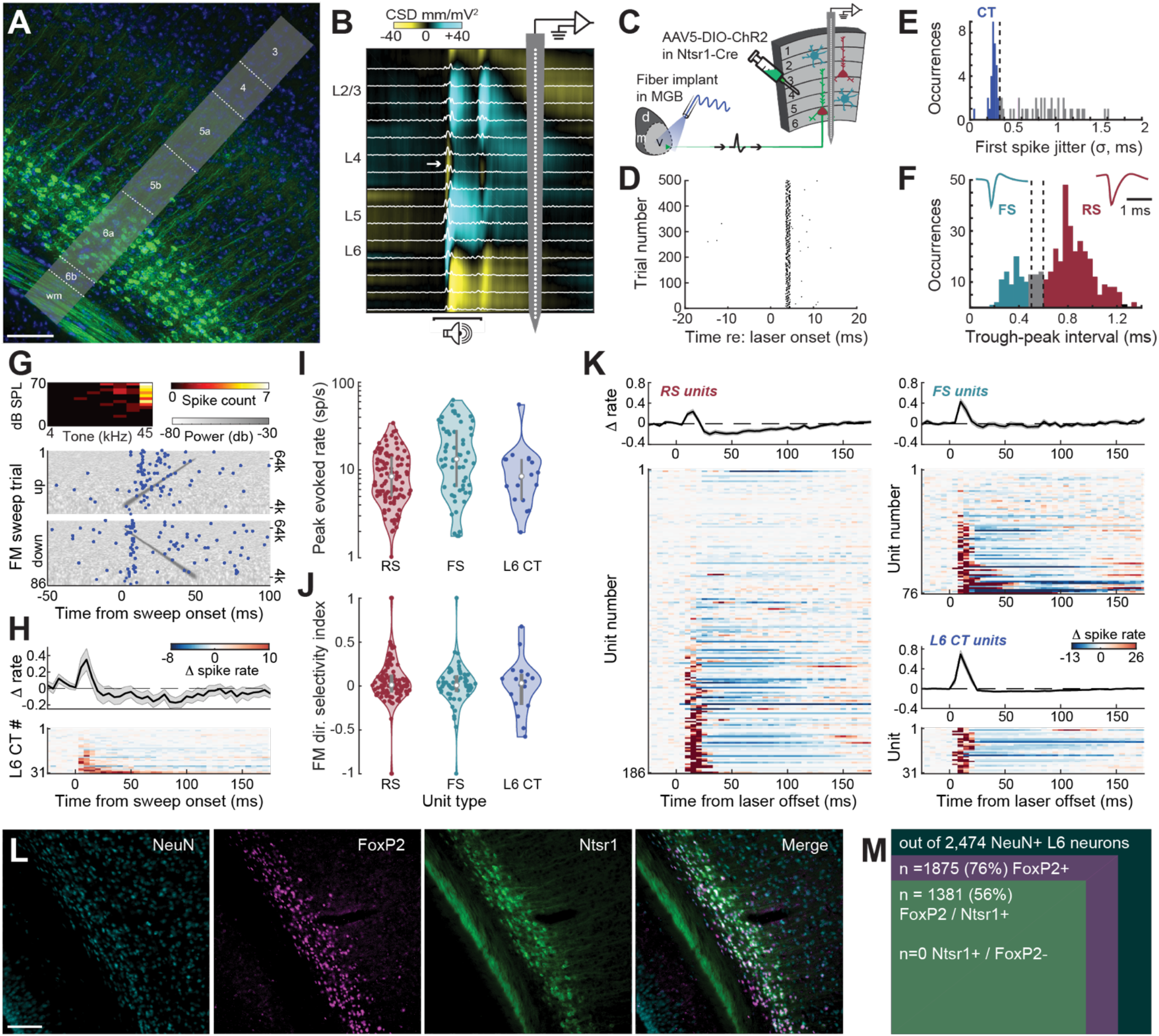
Histological and functional characterization of A1 L6 CT neurons. (**A**) A1 corticothalamic cells labeled in NTSR1-Cre x Ai148 mice have somata in L6a and sparse vertically oriented neuropil labeling up to L5a. Scale bar = 0.1mm. (**B**) Extracellular recordings were made from all layers of A1 with a 64-channel linear probe. Electrophysiological responses are filtered offline to separately analyze unit spiking (white trace) and the current source density (CSD). White arrow in CSD trace identifies the early current sink in layer L4 elicited by a 50ms white noise burst used to assign units to layers. (**C**) Schematic for antidromic optogenetic phototagging in NTSR1-Cre+ L6 corticothalamic neurons that express ChR2. (**D**) Rastergram of antidromic spikes in an example unit elicited by photoactivation of thalamic axon terminals of a L6 corticothalamic unit. (**E**) Histogram of first spike latency jitter in 77 units with optogenetically evoked responses. Directly activated L6 corticothalamic cells were distinguished from indirectly activated units as having a jitter in first spike latency less than 0.35ms (dashed vertical line). (**F**) Isolated single units were classified as FS or RS (trough-to-peak delay < 0.5ms or > 0.6ms, dashed vertical lines). Spike waveforms reflect mean FS and RS waveforms. (**G**) Sensory tuning from an example L6 CT unit. *Top*: Brief tone pips identified a high-frequency receptive field. *Bottom*: FM sweep-evoked spiking rastergrams are superimposed on spectrograms of upward and downward FM sweeps in continuous white noise. (**H**) *Top*: Mean ± SEM normalized firing rates (*top*) and individual unit firing rates *(bottom)* relative to a pre-stimulus baseline for all phototagged L6 CT units (n = 31). (**I**) Peak firing rates elicited by the average of up and down FM sweeps in 108/60/18 significantly responsive RS/FS/L6 CT single units. Firing rates varied significantly between cell types (one-way ANOVA, F = 7.83, p = 0.0005). Post-hoc pairwise comparisons confirmed similar firing rates in L6 CT and other RS units (p = 1). Normalized histogram violin plots present the median value as the white circle and the interquartile range as the grey vertical bars. (**J**) FM direction selectivity index ([up – down] / [up + down]) is similar across sound-responsive cell types (one-way ANOVA, F = 0.6, p = 0.55). (**K**) Neurograms grouped by cell type and present the firing rate change relative to baseline before and after photoactivation of L6 CT units with a 1ms pulse of light. Units are sorted by the absolute value of their mean activity. Line plots present the mean ± SEM normalized firing rates relative to a pre-stimulus baseline for RS, FS and L6 CT cell types. (**L**) A1 cells in NTSR1-Cre x Ai148 mice were immunolabeled for NeuN and FoxP2. Scale bar = 0.1mm. (**M**) Venn diagram depicts the co-localization of FoxP2 in A1 L6 neurons and NTSR1-Cre in FoxP2+ and FoxP2-L6 neurons.

Optogenetics provides a means to artificially manipulate activity in genetically targeted cell types, but can also be used to antidromically identify or “phototag” targeted cell types during extracellular single unit recordings, enabling analysis of cell type-specific activity profiles in response to sensory stimuli, movement or other task-related variables (Guo et al., 2019; Li et al., 2015; Lima et al., 2009; Williamson and Polley, 2019). Here, we used high-density 64-channel linear probes to make extracellular recordings from single units in all layers of A1 in awake, head-fixed Ntsr1-cre mice (**Fig. 1B**). Several weeks prior, we expressed channelrhodopsin-2 in Ntsr1-Cre+ L6 CT neurons and implanted an optic fiber such that the fiber tip terminated above the dorsal surface of the MGB (**Fig. 1C**). This allowed us to optogenetically activate the thalamic axon terminals of L6 CT neurons with a brief (1ms) flash of blue light and quantify the temporal jitter of antidromic spikes recorded from A1 cell bodies (**Fig. 1D**). As confirmed in our recent study, units with minimal trial-to-trial jitter in first spike latency were operationally defined as Ntsr1+ CT cells directly activated by the laser pulse (**Fig. 1E**) (Williamson and Polley, 2019). Units that did not respond to the laser or responded with higher first spike jitter (>0.35 ms) were classified by their spike waveform shape as regular spiking (RS) or putative parvalbumin-expressing GABAergic fast-spiking (FS) neurons (**Fig. 1F**).

We probed the sensory tuning properties of L6 CTs alongside RS and FS units in the same column using a combination of pure tones and frequency modulated (FM) sweeps in a background of continuous white noise (**Fig. 1G**). A subset of optogenetically tagged L6 CT units (n = 18/31 significantly responsive/recorded) exhibited brisk, short-latency responses to FM sweeps (**Fig. 1H**). FM-evoked responses in L6 CTs were not significantly different than other RS units (n = 108/259) or FS units (n = 60/93), either in terms of evoked firing rates (**Fig. 1I**) or FM direction selectivity (**Fig. 1J**, statistical reporting provided in figure legends). The clearest distinguishing feature of L6 CTs was not their sensory response profiles, but rather that photoactivating their axons with just a 1ms pulse of blue light was sufficient to produce an initial wave of intense excitatory responses followed by a prolonged period of suppression in neighboring RS and FS units (**Fig. 1K**). As we have reported previously, sustained periods of local response modulation after laser offset are not observed with photoactivation of other types of A1 corticofugal projection neurons, interneurons or neuromodulatory inputs to A1 (Guo et al., 2017; Williamson and Polley, 2019).

What type of endogenously occurring extrasensory input might play the role of artificial photoactivation, to naturally recruit L6 CT neurons prior to the onset of anticipated sounds? Based on histological characterizations in the visual cortex, we immunolabeled A1 sections from Ntsr1-Cre × Ai148 mice with a neuron-specific marker (NeuN) and for the forkhead box protein P2 (FoxP2), and quantified co-expression between all three markers in L6 (Sundberg et al., 2018). FoxP2 is highly expressed in motor control nuclei, particularly in brain regions supporting audiomotor integration during speech production and vocal motor learning (French and Fisher, 2014). *FOXP2* mutations are linked to vocal communication disorders in humans (Lai et al., 2001), birds (Haesler et al., 2007) and to more generally disrupted audiomotor learning and sensorimotor integration in mice (Castellucci et al., 2016; Kurt et al., 2012). We found that FoxP2 is highly expressed in A1, but only in L6 (**Fig. 1L**). We observed that 74% of L6 FoxP2 neurons in A1 (1381/1875) are Ntsr1-Cre+ and – strikingly – every Ntsr1-Cre+ neuron expressed FoxP2 (**Fig. 1M**). Based on this finding, we hypothesized that movement – or internal motor corollary inputs that precede movement – could be a naturally occurring source of extra-sensory input for L6 CTs.

## Characterizing a highly stereotyped, self-initiated, rapid-onset movement

To characterize the degree and timing of L6 CT activation before and during movement, we needed mice to spontaneously, rapidly and repeatedly deploy a highly stereotyped movement. To this end, we used a minimalistic lickspout sampling task in which tongue contact on a lickspout had a 50% probability of generating a FM sweep after a 0.1s delay and a 50% probability of resulting in a water reward (each independently determined). Sound onset was delayed by 0.1s following lick bout initiation. Continuous background masking noise obscured sound arising from the lick itself. This task design motivated mice to perform sufficient numbers of lick bouts, while ensuring that sound presentation and reward were unpredictable. This task design also allowed us to confirm that sound-evoked A1 responses were suppressed during licking, consistent with many prior reports of suppressed sound-evoked activity during whole body movements including locomotion and bar pressing (McGinley et al., 2015; Rummell et al., 2016; Schneider et al., 2014, 2018; Zhou et al., 2014) (**Supplemental Fig. 1**).

To quantify the timing of orofacial movements and pupil diameter preceding lick bout onset, we used DeepLabCut, a video analysis method for markerless tracking of body movements based on deep neural networks (n = 12 videography sessions in 4 mice; **Fig. 2A**) (Mathis et al., 2018; Nath et al., 2019). We observed a reproducible sequence of orofacial movements associated with licking both within and between mice. Movement of the whisker pad, nose and lower jaw reliably preceded lickspout contact, whereas changes in pupil diameter and pinna movement lagged behind lick bout onset (**Fig. 2B**). Based on this evidence, we proceeded with recording from A1 single units during the lickspout sampling task, knowing that the earliest overt facial movements preceded lickspout contact by approximately 30ms (**Fig. 2C**).

**Figure 2.**
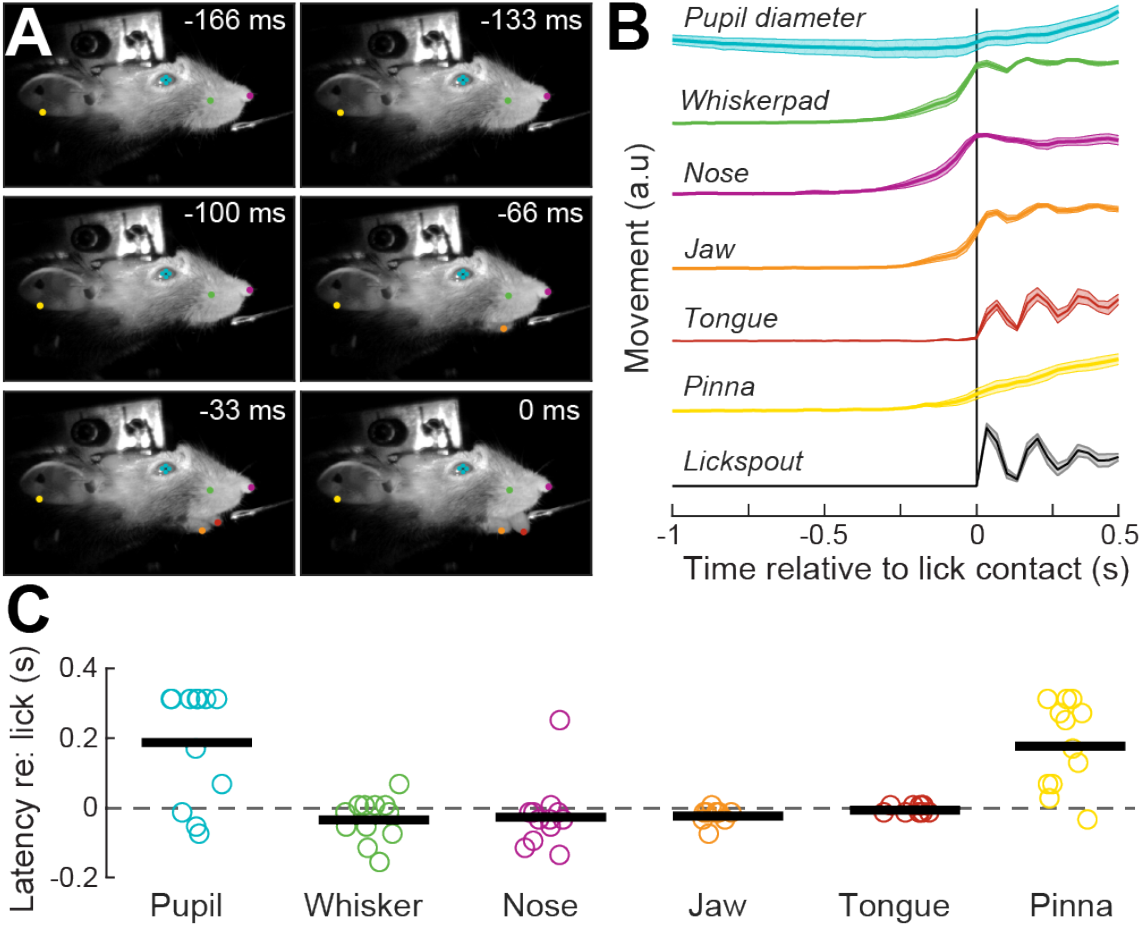
Quantification of a stereotyped, self-initiated, rapid-onset movement sequence culminating in lickspout contact. (**A**) Six consecutive video frames culminating in the moment where the tongue first contacted the lick spout. DeepLabCut marker positions are indicated on each frame for the pinna, whisker pad, nose, tongue, jaw and pupil. (**B**) Mean ± SEM movement amplitudes and pupil diameter are shown relative to electrical contact with the lickspout (n = 12 video sessions from 4 mice). (**C**) Onset latencies for initial changes in facial markers and pupil diameter relative to lickspout contact. Each data point represents a single imaging session. Horizontal bar = sample mean.

## Motor preparatory inputs activate deep layer A1 neurons

Suppression of sound-evoked activity in ACtx RS units during movement is thought to arise, in part, from increased spiking of FS and other local inhibitory neurons prior to movement onset (Schneider et al., 2014). Given the persistent suppressive effect of L6 CT photoactivation on RS units (Fig. 1K), we reasoned that L6 CT units may also be recruited to fire prior to movement onset. As a next step, we characterized the degree and timing of firing rate changes in L6 CT, RS and FS units prior to the onset of self-initiated lick bouts. We found that significantly suppressed spiking during licking was observed in roughly 25% of units and was expressed similarly across layers (n = 28/43/132/135 units in L23/L4/L5/L6, respectively; **Fig. 3A**). Firing rates were significantly increased during licking in approximately 40% of single units, particularly in L6, where spiking ramped up hundreds of milliseconds before contact with the lickspout and was significantly greater than other layers. Averaging over all functional response types, we confirmed prior reports that FS spike rates increased over a 1s period prior to movement onset without any systematic differences across layers (**Fig. 3B, top**) (Rummell et al., 2016; Schneider et al., 2014, 2018). Although a minority of RS units were strongly suppressed prior to lick onset, the overall effect was an increase in RS firing rate that was mainly attributable to strong motor-related activation of L6 units (n = 236, **Fig. 3B, bottom**).

**Figure 3.**
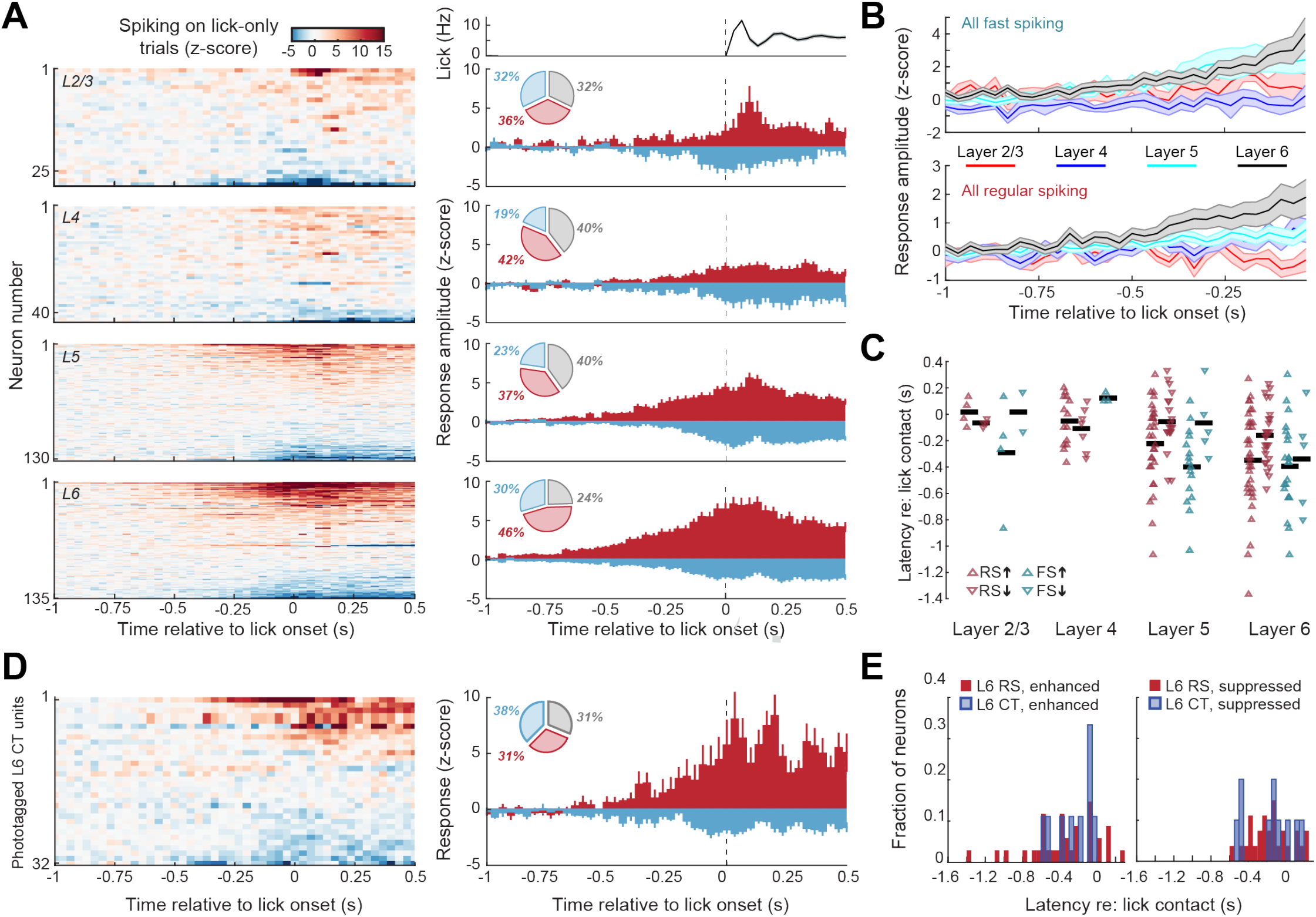
Motor preparatory inputs activate L6 neurons in A1. (**A**) *Left*: Neurograms are grouped by layer and present normalized spike rates for all single units on lick-only trials. Units are sorted by mean normalized activity. *Right*: Pie charts present percentage of units significantly enhanced (red) or suppressed (blue) during the peri-lick period. PSTHs present mean ± SEM normalized firing rates for enhanced and suppressed units from the corresponding layer. Top panel presents the mean ± SEM lick rate during the recording sessions. Firing rate changes among significantly suppressed units were not different across layers (n = 28/43/132/135 units in L23/L4/L5/L6; ANOVA, main effect for layer F = 1.19, p = 0.7). Firing rate changes amongst enhanced units differed between layers and was greatest in L6 (ANOVA, main effect for layer, F = 8.55, p < 0.00003; post-hoc pairwise comparisons with Holm-Bonferroni correction, p = 0.02, 0.00009 and 0.02 for L6 vs L5, L4 and L2/3, respectively). (**B**) Mean ± SEM changes in activity for all FS (top) and RS (bottom) units. *Top*: Firing rate changes in FS units increased prior to movement but was not different across layers (n = 79, Mixed design ANOVA, main effect for time and layer, F = 4.03 and 2.06, p < 1 × 10^−6^ and 0.11, respectively; time × layer interaction term, F = 0.93, p = 0.65). *Bottom*: RS unit activity increased overall prior to lick onset (main effect for time, F = 4.29, p < 1 × 10^−6^), which was mainly attributable to strong motor-related activation of L6 RS units (n = 236, main effect for layer, F = 2.66, p = 0.04; time × layer interaction term, F = 1.58, p < 0.001). (**C**) Motor-related response latencies for each significantly enhanced or suppressed single unit. Horizontal bar = sample mean. (**D**) Motor-related recruitment of optogenetically identified L6 CT units. Neurogram, pie chart and histogram plotting conventions match *A*. (**E**) Histograms of motor-related response latencies for all significantly enhanced (top) or suppressed (bottom) L6 CT units relative to other unidentified enhanced and suppressed RS L6 units.

To determine the latency of motor-related spiking, we followed the analysis approach of prior reports and replicated the finding that FS units activated by movement first begin to increase their spiking hundreds of milliseconds prior to movement onset (mean latency for L2/3, L5, L6 FS units = −292ms, −400ms, −396ms; **Fig. 3C**) (Schneider et al., 2014). Motor-related changes in RS unit spiking tended to lag behind FS units with one notable exception: L6 RS units began to increase their spiking at approximately the same latency as FS units (−349ms prior to lick spout contact; unpaired t-test for enhanced L6 RS vs. FS latencies, p = 0.63). As the earliest changes in L6 RS firing rates begin more than an order of magnitude earlier than the beginning of orofacial movements, they reflect motor preparatory-related activity rather than sensory feedback arising from ongoing movements.

To determine if motor-preparatory recruitment of L6 RS units included L6 CTs, we separately analyzed motor-related activity changes in optogenetically phototagged neurons (n = 32, N = 6 mice; **Fig. 3D**). Latencies for motor related enhancement or suppression were not significantly different than unidentified L6 RS units, which is unsurprising given that the majority of L6 RS units are CT neurons (two-sample K-S test on L6 CT vs L6 RS latency distributions, p > 0.68 for both enhanced and suppressed units; **Fig. 3E**). These results demonstrate that A1 L6 CTs (and possibly other sub-types of deep-layer RS units) begin to ramp up their spiking in response to motor corollary inputs, hundreds of milliseconds prior to movement onset.

## Motor-related inputs from orofacial movements – but not locomotion – recruit A1 L6 corticothalamic neurons

Single unit recordings measure spiking at millisecond resolution, which is useful when calculating onset latencies for neural changes relative to overt movements. However, the yield in single unit recording experiments is much lower than 2-photon imaging, particularly with antidromic phototagging, where we optogenetically tagged less than one CT unit on average in any given recording session. Although the temporal dynamics and relationship to spike coupling with calcium indicators preclude any strong claims about the relative timing of neural activity and movement, we extended our motor characterization experiments with 2-photon imaging to simultaneously visualize larger neural ensembles (**Fig. 4A**).

**Figure 4.**
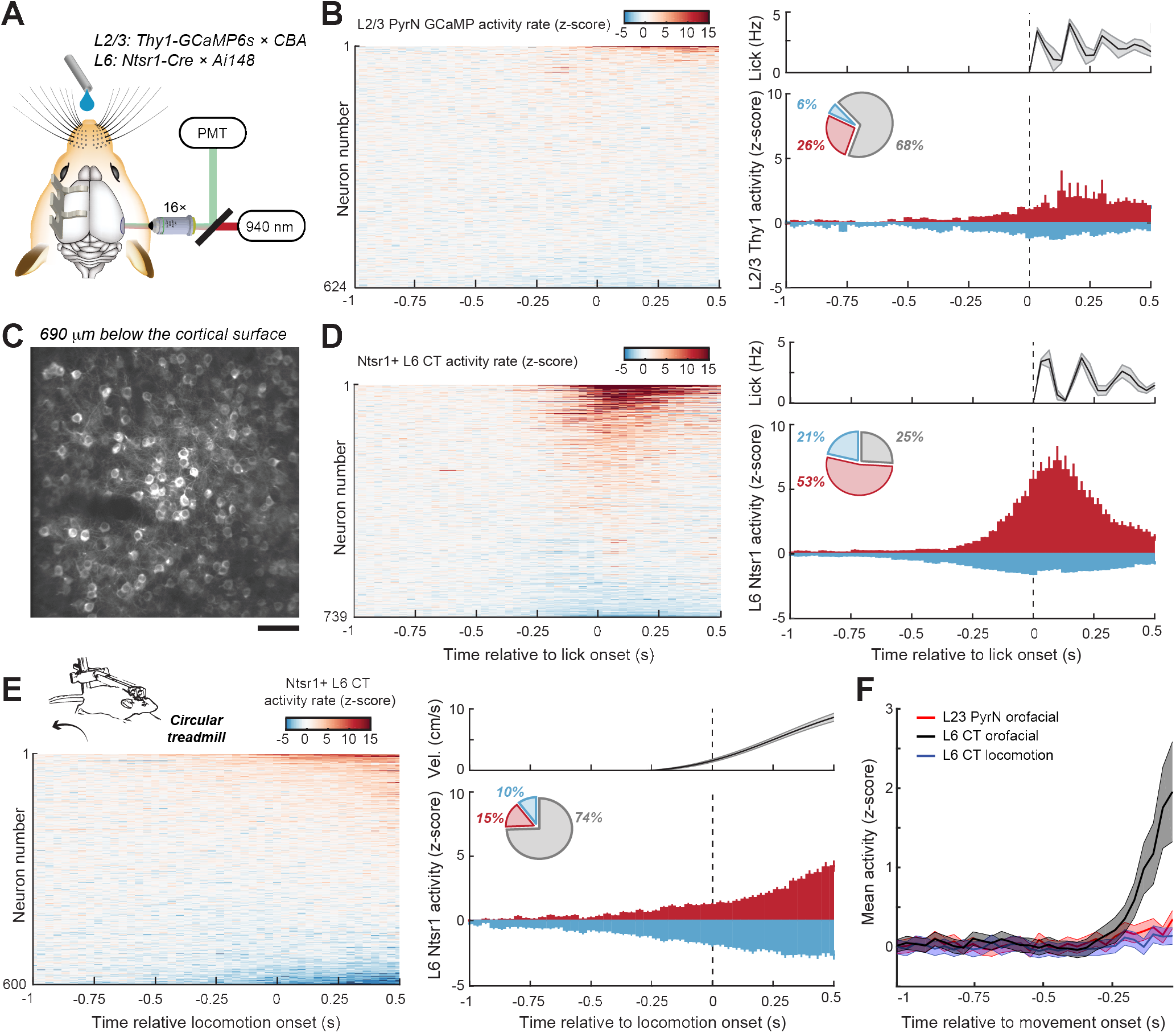
Ensembles of L6 CT neurons are activated by upcoming orofacial movements, but not locomotion bouts. (**A**) Schematic for 2-photon imaging of motor-related activation in L2/3 PyrNs and L6 corticothalamic neurons. (**B**) Lick-related calcium activity from deconvolved GCaMP6 signals in L2/3 PyrNs. *Left*: Neurograms present normalized activity rates for all L2/3 PyrNs on lick-only trials. Neurons are sorted by mean normalized activity. *Right*: Pie charts present percentage of units significantly enhanced (red) or suppressed (blue) during the peri-lick period. PSTHs present mean ± SEM normalized activity rates. Top panel presents the mean ± SEM lick rate for the corresponding recordings sessions. (**C**) GCaMP6 fluorescence in L6 corticothalamic cells in a NTSR1-Cre × Ai148 mouse. Scale bar = 50 μm. (**D**) Lick-related calcium activity in L6 corticothalamic neurons. Graphing conventions match *B.* (**E**) Locomotion-related calcium activity in L6 corticothalamic neurons. Graphing conventions match *B.* Top panel presents the mean ± SEM treadmill velocity from the corresponding recordings sessions. (**F**) Mean ± SEM motor-related changes in PyrN activity for all 2-p imaging experiments. Activity was increased prior to movement onset, but only in L6 CTs and only for orofacial movements related to licking, not locomotion (Mixed design ANOVA, Time × Group interaction, F = 15.49, p = 2 × 10^−144^).

Consistent with our small sample of L2/3 single unit recordings (Fig. 3B-C), orofacial movement elicited weak responses in L2/3 excitatory pyramidal neurons (PyrNs), with 68% of cells not significantly affected by movement, and the remainder of cells mostly activated or suppressed after the lick bout had begun (n = 624; N=5; **Fig. 4B**). Two-photon imaging of L6 cell bodies is impossible under most circumstances on account of fluorescence from superficial layers that greatly reduces the signal-to-noise ratio of somatic GCaMP signals. Because GCaMP expression is limited to L6a in Ntsr1-Cre x Ai148 mice and because these CT cells have sparse, slender, and short apical dendritic and axonal fields (Fig. 1A), there is very little out of plane fluorescence, permitting 2-photon calcium imaging at depths of ~0.7mm without using high excitation laser power (Daigle et al., 2018; Liang et al., 2019; Voigts et al., 2019) (**Fig. 4C**). In agreement with our phototagged recordings, we found that L6 CT neurons were strongly recruited by orofacial movements, where the activity rates of 74% of cells was significantly changed from baseline, mostly reflecting enhanced activity beginning prior to lickspout contact (n = 739; N = 3, **Fig. 4D**).

The most obvious difference between our paradigm and previous work on motor corollary suppression of ACtx RS units was that we focused on orofacial movements, whereas previous studies have used locomotion on a treadmill or other whole body movements (Rummell et al., 2016; Schneider et al., 2014, 2018). To determine if the strong motor-related activation of L6 excitatory neurons reported here could be accounted for based on the type of movement, we performed an additional 2-photon imaging experiment on ensembles of Ntsr1+ L6 CT neurons when mice initiated bouts of running on a treadmill. To our surprise, L6 CT activation prior to movement onset was virtually absent during treadmill running. Whereas 53% of L6 CT neurons showed significantly enhanced activity rates related to orofacial movement, only 15% showed significantly elevated activity around bouts of running (n = 606; N = 3, **Fig. 4E**). In sum, L6 CT PyrNs showed similar patterns of motor-related activity as their L2/3 PyrN counterparts when the source of movement changed from the face to the body (**Fig. 4F**).

## DISCUSSION

### Sources of motor-related inputs to the auditory cortex

Here, we reported the first targeted recordings from L6 CT neurons in a behaving animal. The major discovery was that many L6 CTs and other sub-populations of deep layer putative PyrNs in A1 essentially acted like motor neurons, in that their firing rates increased hundreds of milliseconds prior to movement onset. The sources of movement-related input onto L6 CTs have yet to be determined. Cholinergic projections from the caudal substantia innominata make monosynaptic connections onto ACtx inhibitory neurons to suppress cortical sound processing during states of movement, auditory learning or active listening, though these inputs tend to lag (rather than lead) movement initiation (Abs et al., 2018; Kuchibhotla et al., 2017; Nelson and Mooney, 2016). The best-defined pathway for transmitting movement-related inputs into ACtx comes from neurons in secondary motor cortex (M2) that make monosynaptic connections onto local inhibitory neurons to suppress PyrN responses prior to the initiation of self-generated sounds (Nelson et al., 2013; Schneider et al., 2014, 2018). In addition to their monosynaptic inputs onto local GABA cells, M2 afferents also innervate deep layers of ACtx and reliably elicit excitatory synaptic currents in L6 PyrNs (Nelson et al., 2013), though it has not yet been determined if these cells are L6 CTs and whether M2 input elicits net enhanced spiking rather than suppression in L6 RS neurons. A third possibility is the globus pallidus (GP), based on many reports that GP firing rates are modulated before self-initiated movements (Dodson et al., 2015; Lee and Assad, 2003). The external GP borders the ACtx-projecting regions of substantia innominata and contains a sub-compartment that projects to neocortex, including targeting of deep layer neurons in primary somatosensory cortex (Abecassis et al., 2020; Saunders et al., 2015). In rats, neurons in the external GP and caudate/putamen project to ACtx, though the cell type-specific targets and the balance of projections to A1 versus other cortical fields has not yet been resolved (Chavez and Zaborszky, 2017). One possibility is that there are multiple movement-related pathways that converge on distinct local circuit architectures within A1 to produce distinct forms of cortical response modulation prior to reafferent auditory inputs generated by movement. The precise taxonomy of movements that favor different combinations of motor-related inputs sources and distinct local circuit modulation could be determined in future studies through a comprehensive analysis of self-generated movements alongside inactivation and targeted recordings of key motor systems and local A1 cell types.

### Speculation on a possible role of L6 CT neurons in active listening

Our earlier work noted that optogenetic activation of L6 CTs reset the phase of delta-theta oscillations, modified sensory tuning in the thalamus and cortex, and switched perception into modes of enhanced detection or enhanced discrimination (Guo et al., 2017). Because the largest and clearest effects of L6 CT activation on thalamocortical tuning and perception occurred after photoactivated volleys of L6 spiking had ended, we speculated that L6 CT neurons might be naturally recruited to spike prior to anticipated sounds during active sensing. Here, we discovered that motor corollary inputs may fulfill this role by activating L6 CT neurons prior to sounds generated by orofacial movements. Preparatory facial movements provide a critical reference frame for anticipating the timing of forthcoming vocalizations, provided there are neurons in the auditory system that can both register these motor triggers and also coordinate local circuits and ongoing rhythms to process predicted sound features (Morillon et al., 2015). L6 CTs fulfill both of these requirements, raising the possibility that they might be integrated into distributed motor control systems that support active sensing during vocal motor communication in mice (Okobi et al., 2019). Given the direct linkage between *FOXP2* mutations and monogenic speech and language disorders, one could further speculate that disruptions of auditory L6 CT processing could have devastating consequences on integrating the timing of vocal motor signals with auditory feedback.

## Acknowledgments

We thank Evan Foss and Ishmael Stefanov for contributing to behavioral hardware development and Eyal Kimchi for guidance on using DeepLabCut; We thank D. Kim and the Genetically Encoded Neuronal Indicator and Effector Project at the HHMI’s Janelia Farm Research Campus for making the Thy1-GCaMP6s mouse publicly available. AAV5-Ef1a-DIO hChR2(E123T/T159C)-EYFP was developed by Karl Deisseroth.

## Author contributions

KCC and RSW performed all experiments and analyses on live animals with supervisory input from DBP. TH performed imaging and analysis of fixed tissue. KEH developed neurobehavioral software control; DBP and KKC wrote the manuscript with feedback from all authors;

## STAR ★ Methods

### LEAD CONTACT AND MATERIALS AVAILABILITY

Further information and requests should be directed to and will be fulfilled by the lead contact, Daniel Polley. This study did not generate new reagents.

### KEY RESOURCES TABLE

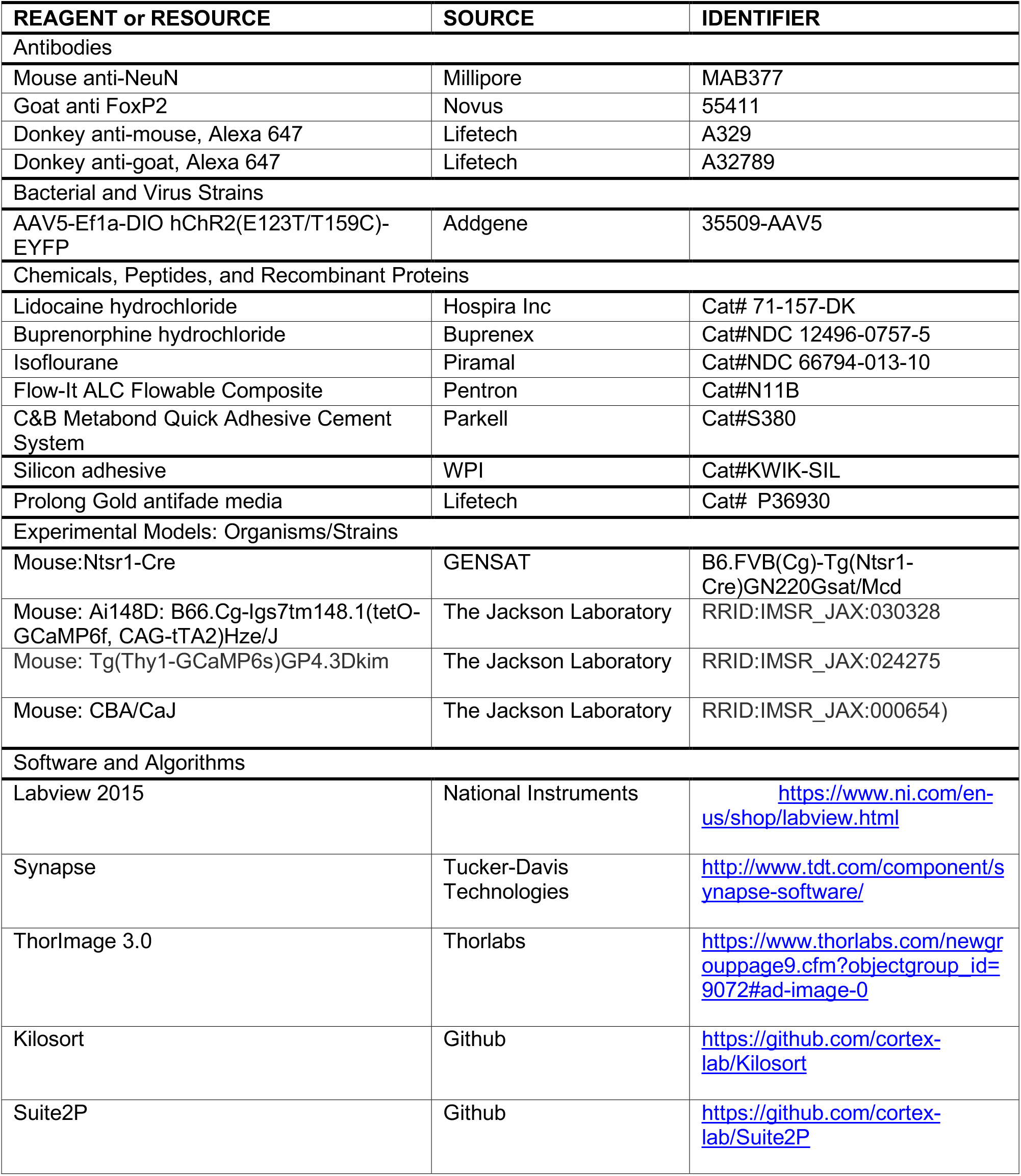
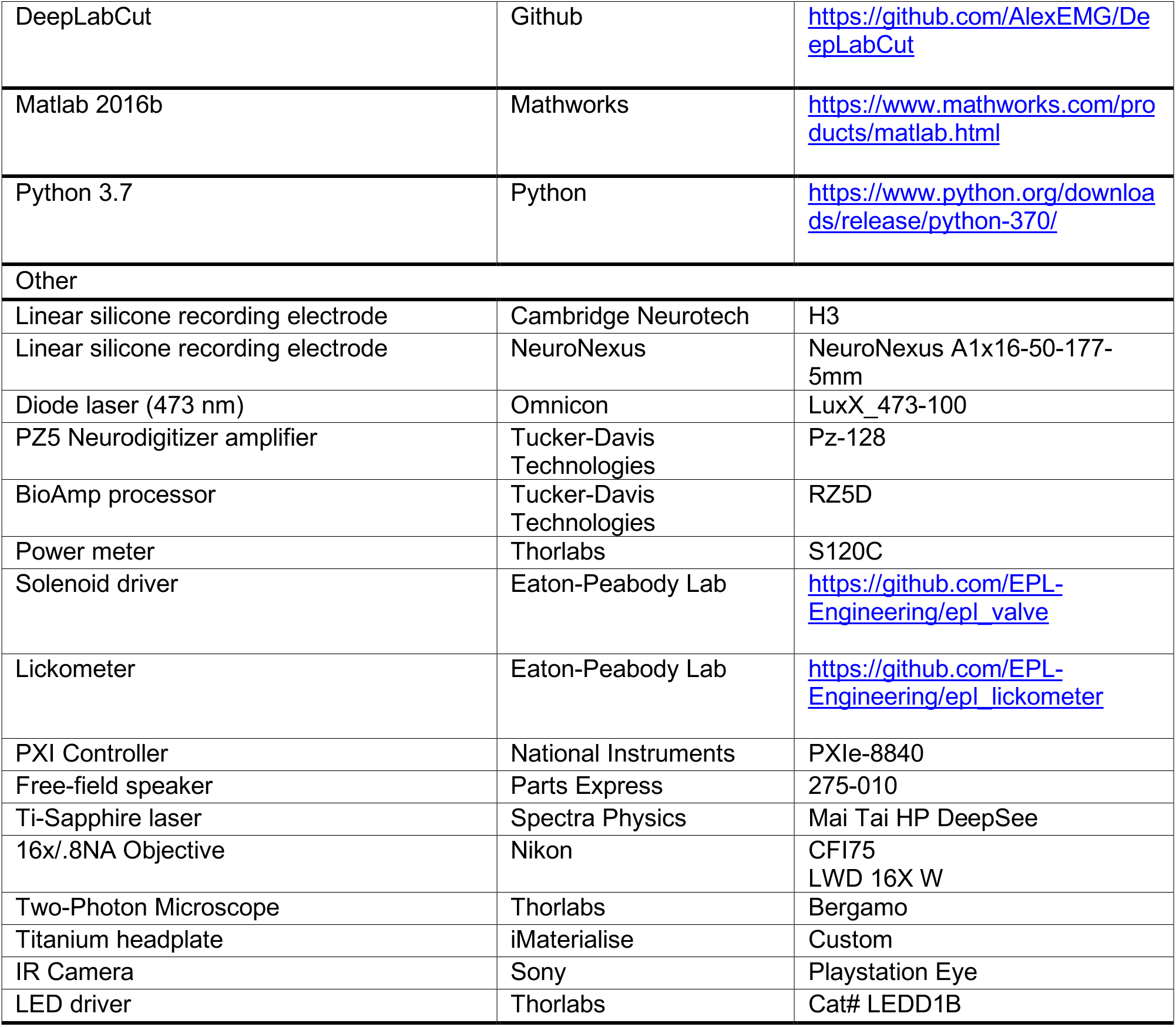

### EXPERIMENTAL MODEL AND SUBJECT DETAILS

All procedures were approved by the Massachusetts Eye and Ear Animal Care and Use Committee and followed the guidelines established by the National Institutes of Health for the care and use of laboratory animals. Mice of both sexes were used for this study. All mice were maintained on a 12 hr light/12 h dark cycle with ad libitum access to food and water, unless undergoing behavioral testing. Mice undergoing behavioral testing were kept on a reversed 12hr light (7:00 a.m.-7:00 p.m) /12 hr dark cycle. Mice were grouped housed unless they had undergone a major survival surgery.

For 2-photon imaging experiments, we used 5 Thy1-GCaMP6sxCBA-CaJ mice and 6 NTSR1-Cre x Ai148 mice. For single-unit electrophysiology experiments, we used 7 NTSR1-Cre mice. For anatomy experiments, we used 3 NTSR1-Cre x Ai148 mice.

### METHOD DETAILS

#### Recovery surgeries

Mice were anesthetized with isoflourane in oxygen (5% induction, 2% maintenance). A homeothermic blanket system was used to maintain body temperature at 36.5° (FHC). Lidocaine hydrochloride was administered subcutaneously to numb the scalp. The dorsal surface of the scalp was retracted and the periosteum was removed. For mice to be used in behavior and physiology experiments, the skull surface was prepped with etchant (C&B metabond) and 70% ethanol before affixing a titanium head plate (iMaterialise) to the dorsal surface with dental cement (C&B metabond). At the conclusion of all recovery procedures, Buprenex (.05 mg/kg) and meloxicam (0.1 mg/kg) were administered and the animal was transferred to a warmed recovery chamber.

#### Lick spout sampling task

Three days after headplate surgery, animals were weighed and placed on a water restriction schedule (1 ml per day). During behavioral training, animals were weighed daily to ensure they remained above 80% of their initial weight and examined for signs of dehydration, such as fur tenting. Mice were given supplemental water if they received less than 1 mL during a training session. Mice were head-fixed in a dimly lit, single-walled sound-attenuating booth (Acoustic Systems), with their bodies resting in conductive cradle. The lickspout apparatus consisted of a single spout positioned 1 cm below the animal’s snout using a 3D micromanipulator (Thorlabs). Tongue contact on the lickspout closed an electrical circuit that was digitized (at 400 Hz) and encoded to calculate lick timing. For electrophysiology experiments, an infrared photobeam emitter/detector for was used to avoid electrical artifacts. Acoustic stimuli were delivered through an inverted dome tweeter positioned 10 cm from the animal’s left ear (CUI, CMS0201KLX). Digital and analog signals controlling sound delivery and water reward were controlled by a PXI system with custom software programed in LabVIEW (National Instruments). Free-field stimuli were calibrated before recording using a wideband ultrasonic acoustic sensor (Knowles Acoustics, model SPM0204UD5).

Mice were initially conditioned to lick the spout to receive a 4-6μL bolus of water 0.5s later. Once mice reliably licked to receive reward, we varied the outcome such that 50% of lick bouts triggered a sound and 50% of lick bouts resulted in reward. Sound and reward probability were determined independently. This task design ensured that both sound presentation and reward were unpredictable, and that sound presentation did not predict reward. Sound onset was delayed by 0.1s following lick bout initiation. Rapid frequency modulated sweeps were used in these experiments because they elicited strong A1 neural responses across a wide range of best frequency tuning (50 ms sweeps presented at 70 dB SPL; 4:64 kHz or 64:4 kHz range [80 octaves/s], 5ms raised cosine onset/offset ramps). To avoid photobleaching, 2-photon imaging sessions were limited to 135 trials per day (approximately 30 minutes) by only presenting upward FM sweeps. For electrophysiology and video tracking experiments, 200-500 trials were performed each day and included both upward and downward FM sweeps.

#### Treadmill running

Head-fixed mice were habituated to running on a low-inertia, quiet cylindrical treadmill over 4-6d period before neural recordings began. Locomotion was recorded using an optical rotary encoder. Locomotion signals were downsampled to 30 Hz and filtered using a zero-phase digital filter. Locomotion was operationally defined as periods where running speed exceeded 2 cm/s. In order to be included for analysis, at least 3s of quiescence were required prior a running bout that lasted for a minimum of 1s. Only sessions that included at least 15 locomotion bouts were used in the analysis. Continuous broadband noise (50 dB SPL) was used to mask sounds generated by running.

#### Two-photon calcium imaging

Three round glass coverslips (one 4mm, two 3mm, #1 thickness, Warner Instruments) were etched with piranha solution and bonded into a vertical stack using transparent, UV-cured adhesive (Norland Products, Warner Instruments). Headplate attachment, anesthesia and analgesia follow the procedure listed above. A 3 mm craniotomy was made over right ACtx using a scalpel and the coverslip stack was cemented into the craniotomy (C&B Metabond). Animals recovered for at least 5 days before beginning water restriction for the lickspout sampling task.

An initial widefield epifluorescence imaging session was performed to visualize the tonotopic gradients of the ACtx and identify the position of A1 as described previously (Romero et al., 2019). Two-photon excitation was provided by a Mai-Tai eHP DS Ti:Sapphire-pulsed laser tuned to 940 nm (Spectra-Physics). Imaging was performed with a 16×/0.8NA water-immersion objective (Nikon) from a 512 x 512 pixel field of view at 30Hz with a Bergamo II Galvo-Resonant 8 kHz scanning microscope (Thorlabs). Scanning software (Thorlabs) was synchronized to the stimulus generation hardware (National Instruments) with digital pulse trains. Widefield and 2-photon microscopes were rotated by 50-60 degrees off the vertical axis to obtain images from the lateral aspect of the mouse cortex while the animal was maintained in an upright head position. Imaging was performed in light-tight, sound attenuating chambers. Animals were monitored throughout the experiment to confirm all imaging was performed in the awake condition using modified cameras (PlayStation Eye, Sony) coupled to infrared light sources.

For 2-photon imaging of L2/3 PyrNs, imaging was performed 175-225 μm below the pial surface. The focal plane for L6 imaging was 600-700 μm below the pial surface, which can be accomplished with relatively low excitation power (107-138 mW) because Ntsr1-Cre neurons have sparse apical dendritic fields that produce minimal out of plane excitation (Daigle et al., 2018; Liang et al., 2019; Voigts et al., 2019). The DeepSee precompensation oscillator (Spectra Physics) was adjusted for each imaging session to improve image quality and reduce laser power for L6 imaging.

Fluorescence images were captured at 2x digital zoom. Raw calcium movies were processed using Suite2P, a publicly available analysis pipeline (Pachitariu et al., 2016). Briefly, movies are registered to account for brain motion. Regions of interest are established by clustering neighboring pixels with similar time courses. Manual curation is then performed in the Suite2P GUI to eliminate for low quality or non-somatic ROIs. Spike deconvolution was performed in Suite2P, using the default method based on the OASIS algorithm (Friedrich et al., 2017).

#### Two-photon imaging analysis

Sound-responsive neurons were identified by measuring deconvolved activity during a prestimulus (−133 – 0ms prior to sound onset) and post-stimulus period (33 – 167ms following sound onset) in sound alone trials. Sound responsive neurons were operationally defined as having significantly elevated activity in the post-stimulus period based on a one-tailed paired-test (p < 0.05). To calculate sound-evoked activity rates for sound alone and sound + lick conditions, the baseline activity level during the pre-stimulus window was subtracted from the post-stimulus window.

The modulation index for sound-evoked activity was calculated as:

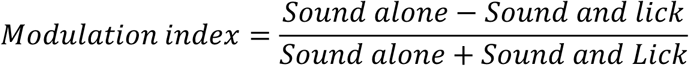

To calculate activity rates during lick only conditions, we first defined baseline activity during a 0.5s period that was separated by at least 2s from the last lick of a preceding bout and 1.5s from the first lick of an upcoming bout. Lick-related activity was calculated during a 1.5s period beginning 1s prior to lick onset, where mean activity in each 33ms time bin was expressed in units of z-score relative to the baseline distribution. Neurons were classified as significantly enhanced or suppressed during licking by calculating the mean activity rates during a period of licking (−0.5 to 0.5s relative to lick onset) versus a period of quiescence (−2 to −1s relative to lick onset) based on a p < 0.05 threshold of a two-tailed Wilcoxon rank-sum test.

#### Virus mediated gene-delivery

For mice used in optogenetics experiments, two burr holes (<1mm diameter each) were made in the skull overlying A1, 1.75 – 2.25mm rostral to the lambdoid suture. A motorized injection system (Stoelting) was used to inject 200 nL of AAV5-Ef1a-DIO hChR2(E123T/T159C)-EYFP into each burr hole 600 μm below the pial surface. Electrophysiology experiments began 3-4 weeks after injection. Immediately following injections, the scalp was sutured and the animal recovered in a warmed chamber. All virus solutions were injected at a rate of 15 nL/min.

#### Preparation for single unit-recordings in head-fixed mice

A ground wire (AgCl) was implanted over the left occipital cortex. A craniotomy was made above the MGB (centered 2.75 mm lateral to the midline and 2.75 mm caudal to bregma) and the position of the ventral subdivision was identified with a cursory mapping of pure tone receptive fields as described previously (Guo et al., 2017; Williamson and Polley, 2019; Williamson et al., 2015). An optic fiber (flat tip, 0.2mm diameter, Thorlabs) was inserted 2.4 – 2.8mm to rest 0.2mm above the physiologically identified dorsal surface of MGB. The fiber assembly was cemented in place and painted with black nail polish to prevent light leakage. Animals were allowed to recover for at least three days before water restriction began.

On the day of the first recording session, mice were briefly anaesthetized with isoflurane (2%) and a small craniotomy (0.5 medial-lateral x 1.25 mm rostral-caudal) was made over A1, centered 2mm from the lambdoid suture. Each day, a small well was made around the craniotomy with UV-cured cement and filled with lubricating ointment (Paralub Vet Ointment). At the conclusion of each recording, the chamber was flushed, filled with new ointment, and capped with UV-cured cement.

#### Single unit recordings in head-fixed mice

A 64-channel silicon probe (H3 probe, Cambridge Neurotech) was slowly advanced (100 μm/min) into ACtx perpendicular to the pial surface until the tip of the electrode was 1.3-1.4mm below the cortical surface, ensuring full coverage of all layers of A1. The brain was allowed to settle for at least 15 minutes before recordings began. On the day of the first recording, multiple penetrations were made to identify the tonotopic reversal which represents the rostral border of A1 (Hackett et al., 2011).

Raw neural data was digitized at 32-bit, 24.4 kHz (Neurodigitizer and RZ5 BioAmp Processor; Tucker-Davis Technologies) and stored in binary format. To eliminate artifacts, the common mode signal (channel-averaged neural traces) was subtracted from all channels in the brain. Signals were notch filtered at 60Hz, then band-pass filtered (300-3000 Hz, second order Butterworth filters). Raw signals were notch filtered at 60 Hz and downsampled to 1000 Hz to extract local field potentials. The CSD was calculated as the second spatial derivative of the LFP signal. To eliminate potential artifacts introduced by impedance mismatching between channels, signals were spatially smoothed along the channels with a triangle filter (5-point Hanning window). Two CSD signatures were used to identify L4 in accordance with prior studies: A brief current sink first occurs approximately 10ms after the onset of broadband noise burst (50ms duration, 70 dB SPL, 100 trials), which was used to determine the lower border of L4. A tri-phasic CSD pattern (sink-source-sink from upper to lower channels) occurs between 20ms and 50ms, where the border between the upper sink and the source was used to define the upper boundary of L4 (Guo et al., 2017; Metherate et al., 2005; Muller-Preuss and Mitzdorf, 1984).

Kilosort was used to sort spikes into single unit clusters (Pachitariu et al., 2016). Single-unit isolation was based on the presence of both a refractory period within the interspike interval histogram, and an isolation distance (>10) indicating that single-unit clusters were well separated from the surrounding noise (Schmitzer-Torbert et al., 2005). RS and FS designation was based on the ratio of the mean trough to peak interval > 0.6 (RS) or < 0.5 (FS). Ntsr1-Cre L6 corticothalamic units were identified using antidromic phototagging. A 1ms pulse of 473nm light was delivered via the implanted MGB fiber to fire antidromic spikes in L6 corticothalamic units that expressed hChR2 (5-45 mW laser power, repeated at 4Hz for 500 repetitions with a diode laser, Omicron, LuxX). Ntsr1+ L6 corticothalamic cells were distinguished from indirectly activated neurons based on laser-evoked spiking rates at least 5 sd above baseline and first spike latencies that varies by less than 0.35 SD, as confirmed in a previous study (Williamson and Polley, 2019).

#### Single unit recording analysis

Single unit spike times were binned at 200 Hz for sensory characterization and laser-evoked firing rate analyses. Sound- and laser-evoked spiking was referenced to the mean spike rate 100ms before stimulus onset. Frequency response areas for sound responsive units were obtained by presenting pseudorandomly ordered pure tones (50 ms duration, 4 ms raised cosine onset/offset ramps) of variable frequency (4–64 kHz in 0.5 octave increments) and level (0–60 dB SPL in 5 dB SPL increments). Each pure tone/level combination was repeated two times. Spikes were collected from the tone-driven portion of the PSTH and averaged across repetitions.

Single unit spike times for lick-related analyses were binned at 30 Hz for direct comparison with 2-photon and behavior tracking data. Units were excluded from further analysis if spontaneous spike rates were < 0.1 Hz. Sound-responsive units, evoked-activity rates and the modulation index for sound alone and sound + lick conditions were calculated similarly to the 2-photon data except that pre- and post-stimulus spike measurement windows were -66-0ms and 33-99ms relative to sound onset, respectively. Lick related activity was calculated with the same methods as described above for 2-photon imaging. Latencies of motor-related excitation or suppression was defined as the first time point that activity consistently exceeded the 5^th^ or 95^th^ percentile of the baseline spike rate distribution (as per Schneider et al, 2014). Analyses was initiated at the minimum (suppressed) or maximum (enhanced) activity time bin and preceded backwards in time until this condition was first met.

#### Capture and quantification of orofacial movements

Videos of the animal’s face were obtained using a camera (Playstation Eye, Sony) coupled with infrared light sources (Thorlabs) and a 5-50 mm varifocal lens (Computar CS-Mount). Videos were acquired at 30 Hz (N=2) or 60 Hz (N=2) at a resolution of 512×512 pixels. A single point on the vibrissae array, nose, jaw, tongue and pinna were labeled alongside four cardinal positions on the circumference of the pupil in 60 frames for each imaging session. Manually labeled frames were split into training and test sets and the network was trained for 1.3 million iterations using default parameters in DeepLabCut (Mathis et al., 2018; Nath et al., 2019). Low-confidence outlier frames were manually re-labeled, the model re-trained and full video dataset reanalyzed. As the lower jaw and tongue markers were not visible until lick initiation, the point where they first appeared was manually entered as their position for all quiescent frames. Any remaining spurious position values where model confidence was < 0.9 was replaced with the interpolated value determined from surrounding frames.

Each tracked point was expressed as a 3-D vector as X × Y × Time. All videos recorded at 60 Hz were downsampled to 30 Hz for further analysis. Pupil diameter was calculated as the mean of the two diameter measurements. All other points were reduced to a 1-dimensional activity trace d(t) as:

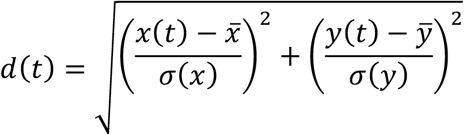

where x(t) is the x position at timepoint t and y(t) is the y position at time point t, which are normalized using the mean and the standard deviation of the entire x and y position traces respectively. A zero phase digital filter was then used to smooth activity traces without introducing temporal distortions. Movement latency was computed identically to neural latencies.

#### Anatomy

Mice were deeply anaesthetized with ketamine and transcardially perfused using 4% paraformaldehyde in 0.01M phosphate buffered saline. Brains were extracted and post-fixed in 4% paraformaldehyde for 12 hours, then placed in 30% sucrose. Brains were sectioned coronally at 40 μm into 0.1M phosphate buffer (PB), preserving all sections containing cerebral cortex. To co-localize Ntsr1-Cre and FoxP2 in NeuN-labled ACtx neurons, fluorescent immunohistochemistry assays were performed on sections of NTSR1 × Ai148 brains. Sections were rinsed in 0.1M phosphate buffered saline (PBS), permeabilized for 2 hours at room temperature (RT) in a blocking solution of 0.1M PBS, 2% BSA, 5% donkey serum, 0.1% Tween-20, then incubated for 48 hours at 4°C in blocking solution containing the primary antibodies. Sections were rinsed in PBS then incubated for 2 hours at RT in blocking solution containing secondary antibodies, counterstained in DAPI for 5 minutes, rinsed in 0.01M PB, mounted onto glass slides, then coverslipped using Prolong Gold antifade media (Lifetech, Eugene OR).

ACtx sections were imaged in z-stacks over 20 um (2 um steps) on a Nikon 90i epifluorescence microscope (40x Plan Fluor, oil DIC N2). Co-localization of NeuN, NTSR1-Cre labeled GFP, and FoxP2 were quantified in 0.25mm wide regions of layer 6 in A1. Labeled cells were counted independently for each fluorescence channel in 2 adjacent sections from each hemisphere of three brains using the Taxonomy function in Nikon Elements AR software. Analysis was restricted to NeuN-labled cells containing a DAPI-labeled nucleus.

### QUANTIFICATION AND STATISTICAL ANALYSIS

All statistical analyses were performed in MATLAB 2016b (Mathworks). Data are reported as mean ± SEM unless otherwise indicated. Non-parametric tests were used when data samples did not meet assumptions for parametric statistical tests. Inflated familywise error rates from multiple comparisons of the same sample were adjusted with the Holm-Bonferroni correction. Statistical significance was defined as p < 0.05.

### DATA AND CODE AVAILABILITY

Data acquisition and analysis was performed with custom scripts in MATLAB, LabVIEW, and Python. Spike sorting was done in Kilosort (https://github.com/cortex-lab/Kilosort). Two photon imaging ROI extraction and spike deconvolution was done in Suite2p (https://github.com/cortex-lab/Suite2P). Markerless behavior tracking was done with DeepLabCut (https://github.com/AlexEMG/DeepLabCut).

### ADDITIONAL RESOURCES

Not applicable.

## SUPPLEMENTAL MATERIALS

**Supplemental Figure 1.**
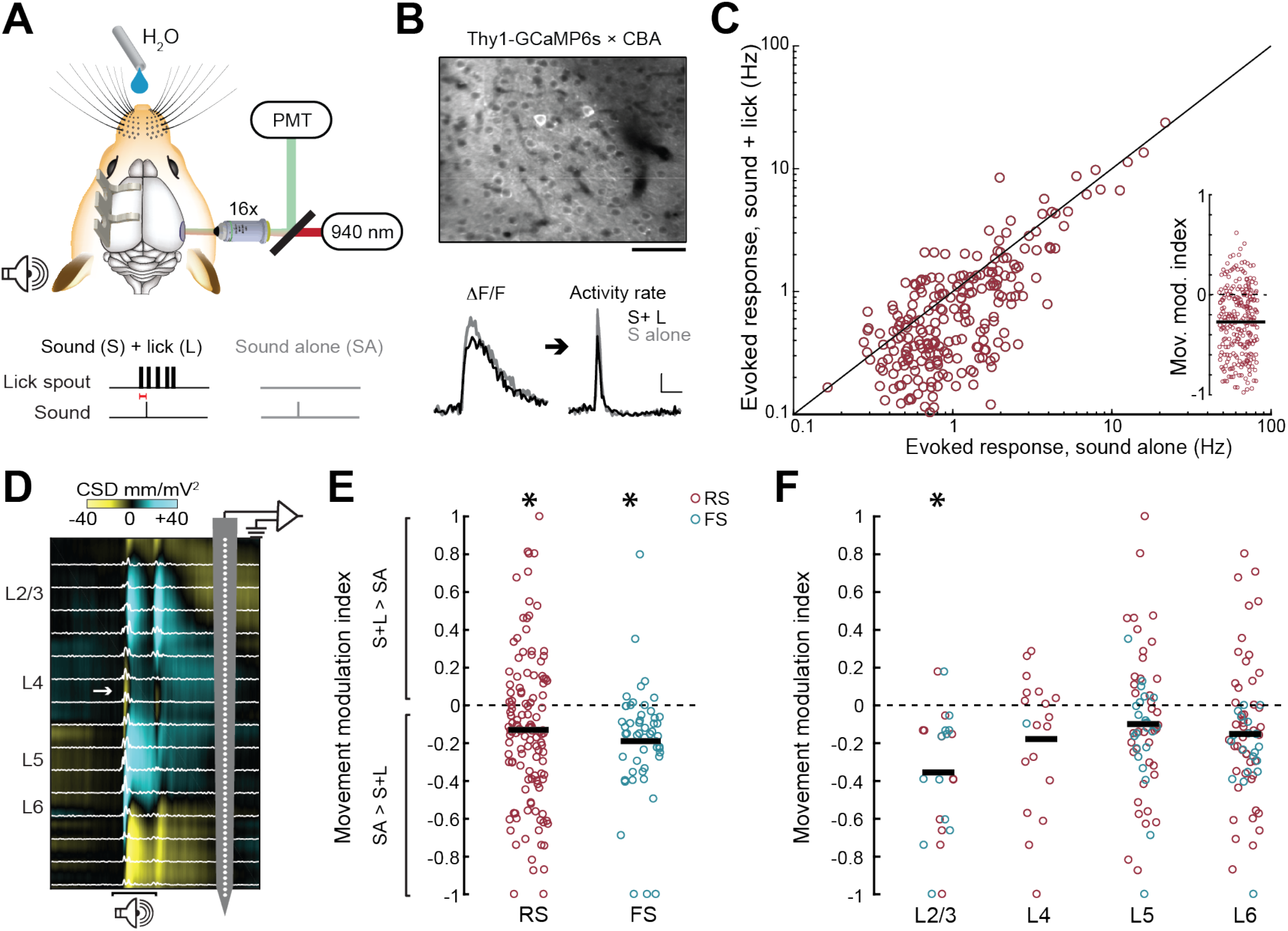
Suppression of lick-triggered sounds in A1. (**A**) 2-photon imaging of A1 L2/3 PyrN activity during a lick spout sampling behavior in which a frequency modulated sweep is either presented 0.1s following lick initiation or during periods of rest. (**B**) *Top*: GCaMP6s fluorescence from L2/3 of a *Thy1GCaMP6s* × CBA mouse. Scale bar = 50 μm. *Bottom*: Sound-evoked fractional change in fluorescence (left) and deconvolved activity rate (right) from a L2/3 PyrN during licking or quiescence. Horizontal scale bar = 0.5s, Vertical scale bar = 20% dF/F (left) and 5 deconvolved events per second (right). (**C**) Suppression of lick-triggered sound responses in L2/3 PyrNs (paired t-test, p < 1 × 10^−6^). *Inset*: The same data presented as a movement modulation index where values > 0 reflect facilitated responses during licking and < 0 reflect suppressed responses during licking. (**D**) Extracellular recordings were made from all layers of A1 with a 64-channel silicon probe during the lick spout sampling behavior. Electrophysiological responses are filtered offline to separately analyze unit spiking (white trace) and the current source density (CSD). White arrow in CSD trace identifies the early current sink in layer L4 elicited by a 50ms white noise burst used to assign units to layers. (**E**) Scatterplots present movement modulation index for every sound-responsive single unit (circle) according to RS or FS spike shape. Significant movement-related suppression was observed both in RS and FS units (one-sample t-test against a population mean of 0, p < 0.0005 for both RS and FS, n = 125 and 52, respectively from 7 mice) with no difference in the degree of suppression between cell types (unpaired t-test for RS vs FS, p = 0.33. Horizontal bars denote sample means. Asterisks indicate p < 0.05 with a one-sample t-test against a population mean of 0 after correcting for multiple comparisons. (**F**) Scatterplots present movement modulation index for every sound-responsive single unit (circle) according to cortical layer. Laminar differences went in the opposite direction of prior work, in that we observed weaker and more variable spiking suppression to self-generated sounds in deeper cortical layers than more superficial layers (Rummell et al., 2016; Schneider et al., 2018).

